# Mobility shift-based electrophoresis coupled with fluorescent detection enables real-time enzyme analysis of carbohydrate sulfatase activity

**DOI:** 10.1101/2020.12.01.406876

**Authors:** Dominic P Byrne, James A London, Patrick A Eyers, Edwin A Yates, Alan Cartmell

**Affiliations:** Department of Biochemistry and Systems Biology, Institute of Systems, Molecular and Integrative Biology, Biosciences Building, Crown Street, University of Liverpool, Liverpool L69 7ZB, United Kingdom

**Keywords:** Carbohydrate sulfatase, inhibitor, mobility-shift, NMR, GlcNAc. Glycosaminoglycan

## Abstract

Sulfated carbohydrate metabolism is a fundamental process, which occurs in all domains of life. Carbohydrate sulfatases are enzymes that remove sulfate groups from carbohydrates and are essential to the depolymerisation of complex polysaccharides. Despite their biological importance, carbohydrate sulfatases are poorly studied and challenges remain in accurately assessing the activity, specificity and kinetic parameters. Most notably, separation of desulfated products from sulfated substrates is currently a time-consuming process. In this paper, we describe the development of rapid capillary electrophoresis coupled to substrate fluorescence detection as a high-throughput and facile means of analysing carbohydrate sulfatase activity. The approach has utility for the determination of both kinetic and inhibition parameters and is based on existing microfluidic technology coupled to a new synthetic fluorescent 6S-GlcNAc carbohydrate substrate. Furthermore, we compare this technique in terms of both time and resources, to high performance anion exchange chromatography and NMR-based methods, which are the two current ‘gold standards’ for enzymatic carbohydrate sulfation analysis. Our study clearly demonstrates the advantages of mobility shift assays for the quantification of near real-time carbohydrate desulfation by purified sulfatases, and could support the search for small molecule inhibitors of these disease-associated enzymes.

**One sentence summary:** Sulfatases remove sulfate groups from biomolecules; in this study we report a rapid and robust capillary electrophoresis assay for the quantification of carbohydrate desulfation.

## Introduction

Sulfated glycans are essential, often complex, molecules present in most eukaryotes and in all metazoans [1, 2]. For instance, the sulfated glycans of the glycosaminoglycan (GAG) class, which include heparan sulfate (HS), chondroitin sulfate (CS), dermatan sulfate (DS) and keratan sulfate (KS), are found ubiquitously in the glycocalyx and extracellular matrix of all mammals [3]. The GAGs regulate the development of metazoan anatomy and organisation, extracellular cell signalling, growth and homeostasis [3–5]. These effects are mediated through interactions with target proteins, whose identity often depends on a combination of shape and charge complementarity. The Protein:GAG binding interaction is driven initially through charge interactions between the negatively charged (carboxylate and multiple sulfate) groups of the GAGs and the positive lysine or arginine (and, if protonated, histidine) side chains of their protein binding partner(s). These ionic interactions are complemented by additional hydrogen bonds and the relative flexibility of the GAGs, both around the glycosidic linkage and, in the case of 2-O sulfated L-iduronic acid (L-IdoA2S), a major constituent of HS and DS, the conformation (^1^C_4_/ ^2^S_0_ equilibrium) of these residues. This enables, in some instances, high affinity (*K*_d_ ~10^-9^ M) binding interactions to occur, as is well-documented for fibroblast growth factors (FGFs)[6]. Interactions with specific protein partners[7, 8] are dependent on both the extent and distribution of sulfate groups in the GAGs and the resulting charge and conformational consequences [9, 10]. Site-specific sulfate groups can be edited enzymatically by a variety of specific sulfatases [11, 12]. In the mammalian extracellular context, there are currently two known sulfatases, termed SULF-1 and SULF-2, which possess distinct HS sequence specificities [13, 14]. Both SULFS act in an *endo*-fashion to remove 6-O-sulfate groups from *N*-acetyl-d-glucosamine (GlcNAc) residues (i.e. from 6S-GlcNAc) of HS. On the other hand, the turnover of GAGs in cells is achieved by *exo*-acting lysosomal enzymes, a series of specific enzymes including glycosidases and sulfatases, whose sequential action degrades the target GAG into monosaccharide units and sulfate anions [15]. Importantly, impairment of these enzymes leads to the accumulation of partial breakdown products, resulting in a series of metabolic disorders termed mucopolysaccharidoses (MPS) [16, 17].

Glycosaminoglycans are also important for the vast microbial community found in the colon, and provide a constant, high priority, source of nutrients [18–20] that are crucial for maintaining the microbiome, which plays a central role in host immunity and metabolism [21–23]. Lining the colonic tract is the mucopolysaccharide mucin, composed mainly of MUC2, a proteoglycan which is 80 % carbohydrate by mass, and heavily sulfated, up to 10 % by mass [24]. MUC2 is an ever present prokaryotic nutrient source, but also an important colonisation factor, and its aberrant desulfation has been linked to chronic inflammatory bowel diseases [21, 23, 25]. Finally, macroalgae cell wall matrix polysaccharides can also be heavily sulfated, a feature which is vital for their function and metabolism. These ‘marine’ glycans, which include fucoidan sulfate and carrageenan, are important industrial resources, valuable as carbon reservoirs, and serve as sources of nutrients for marine bacterial ecosystems [26–28]. Again, de-sulfation is an obligatory step in the metabolism of these molecules by marine bacteria.

De-sulfation of glycans is carried out by carbohydrate sulfatases, which are catalogued in the SulfAtlas database (http://abims.sb-roscoff.fr/sulfatlas/) [29] into four broad families, termed S1-S4. These families share a common fold, mechanism and catalytic apparatus. The S1 family, which contains most of the carbohydrate sulfatases identified to date is further divided up into 72 subfamilies designated S1_X, is found in all domains of life, and family members utilise a non-standard amino acid, formylglycine (FG), as the catalytic nucleophile for sulfate ester bond cleavage. Formylglycine is installed co-translationally into the consensus sequence Cys/Ser-X-Pro/Ala-Ser/X-Arg at the Cys/Ser position [30].

The study of carbohydrate sulfatases poses many significant challenges, most notably, absolute quantification of their enzymatic activities. Sulfated carbohydrates themselves can be difficult to detect, relying on low throughput, expensive and often cumbersome procedures, such as mass spectrometry, NMR and HPLC. High performance anion exchange chromatography (HPAEC) with pulsed amperometric detection (PAD), is currently considered the gold standard for detection of most mono-, di- and oligosaccharides, but does not perform especially well for sulfated molecules. Its use is therefore limited largely to detecting the enzymatic degradation products (*i.e*. un-sulfated sugars) of mono-sulfated carbohydrates. Moreover, detecting the release of the sulfate anion itself is extremely challenging and few quantitative methods exist, especially at the low levels usually required for analytical work with limited reagents. NMR spectroscopy offers a potential solution, but is expensive in terms of time and substrate quantities, as well as requiring expertise and access to specialist equipment. Added to these considerations, the co-translationally-formed formylglycine catalytic residue of S1 family sulfatases, is sometimes incorporated in low abundance in recombinant systems. Overall, these considerations present a challenging set of circumstances for the careful analysis of this critical, but largely understudied, class of carbohydrate-active desulfation enzymes.

We previously described an *in vitro* method for assaying heparan sulfate 2-*O*-sulfotransferase (HS2ST)-driven sulfation of a hexasaccharide substrate using a capillary electrophoresis (CE)-based system coupled to fluorescence detection [31]. In this approach, the enzymatic transfer of the charged sulfate unit from the 3’-phosphoadenosine 5’-phosphosulfate (PAPS)-nucleotide cofactor to the 2-O position of the heparan sulfate IdoA residue leads to a change in the physicochemical properties (in this case an increase in negative charge) of the resulting product compared to the fluorescently tagged (fluorescein) substrate, which can be detected as a real-time mobility shift in a high-throughput microfluidic assay format. This technique was originally exploited for the analysis of peptide phosphorylation by protein kinases, although it can also be employed to detect peptide de-phosphorylation by protein phosphatases [32, 33]. Moreover, in principle, the addition or removal of any modification on a fluorescent substrate that can be detected through a charge-based mobility shift is now feasible. Here, we describe a novel use of this technology to develop a highly sensitive, quantitative, high-throughput and user-friendly technique to study carbohydrate de-sulfation by a purified recombinant sulfatase *in vitro*. We compare the efficiency of this technique to HPAEC and NMR methods, by employing the S1_11 sulfatase termed BT4656, and the 6-O-sulfated substrate 6S-GlcNAc [20], labelled with the fluorophore, BODIPY-FL hydrazide, to illustrate the utility of the assay for the cheap and rapid kinetic analysis of carbohydrate sulfatases, and their real-time inhibition by substrate-mimetics.

## Results and discussion

### Comparison of capillary electrophoresis (CE), HPAEC and NMR methods for the determination of carbohydrate sulfatase kinetics

We studied the enzymatic cleavage of O6-linked sulfate from a novel fluorescently labelled 6S-*N*-acetyl-d-glucosamine (6S-GlcNAc-BODIPY) substrate (**Figure 1A**) by the highly purified recombinant [Bacteroides thetaiotaomicron](https://www.rcsb.org/search?q=rcsb_entity_source_organism.ncbi_scientific_name:Bacteroides%20thetaiotaomicron%20VPI-5482) sulfatase BT4656^S1_11^, in which Ser 77 was substituted with Cys prior to expression in *E. coli;* this mutation is required for *E.coli* to generate the formylglycine nucleophile (**Figure 1B**), while the wild-type sulfatase with Ser 77 in the active site is catalytically inactive, but can bind to substrates (Figure 1C). Specific binding to the cognate substrate 6S-*N*-acetyl-glucosamine, but not the epimer 6S-*N*-acetyl-galactosamine, was confirmed using a previously-validated DSF binding assay [31] (**Figure 1C**). We next evaluated a new CE-based microfluidic sulfatase assay that detects real-time changes in substrate structure (in this case, specific removal of a negatively charged sulfate group) through application of an electric field. This enabled the real-time, ratiometric quantification of sulfate hydrolysis in solution, by comparing the change in retention time of the separated sulfated substrate (6S-GlcNAc-BODIPY) and de-sulfated product (GlcNAc-BODIPY) of the enzyme assay. The fluorescent label 4,4-difluoro-5,7-dimethyl-4-bora-3a,4a-diaza-*s*-indacene-3-propionic Acid, hydrazide (BODIPY-FL hydrazide) was selected (see below), despite a relatively high initial cost, owing to its high extinction coefficient (80,000 M^-1^cm^-1^) and its stability over a variety of pH values (pH 3 – 12) that is critical for analysis in buffer-sensitive assays [34]. The labelled substrate was detected in a 20 nl volume in-line *via* LED-induced fluorescence, resulting in a time-dependent leading sulfated peak and a smaller secondary peak that represented the initial non-sulfated substrate (**Figure 2A**). The fluorescent intensity (measured as both peak height and differentiated area of the peak) increased in a linear fashion with 6S-GlcNAc-BODIPY substrate concentrations ranging between 0.25 and 15 μM (**Figure 1C, D**). Above this range, linearity deteriorated due to saturation of the fluorescence signal. As such, analysis of enzymatic rates of de-sulfation are less accurate at concentrations above 15 μM for BODIPY-labelled substrates.

**Figure 1.**
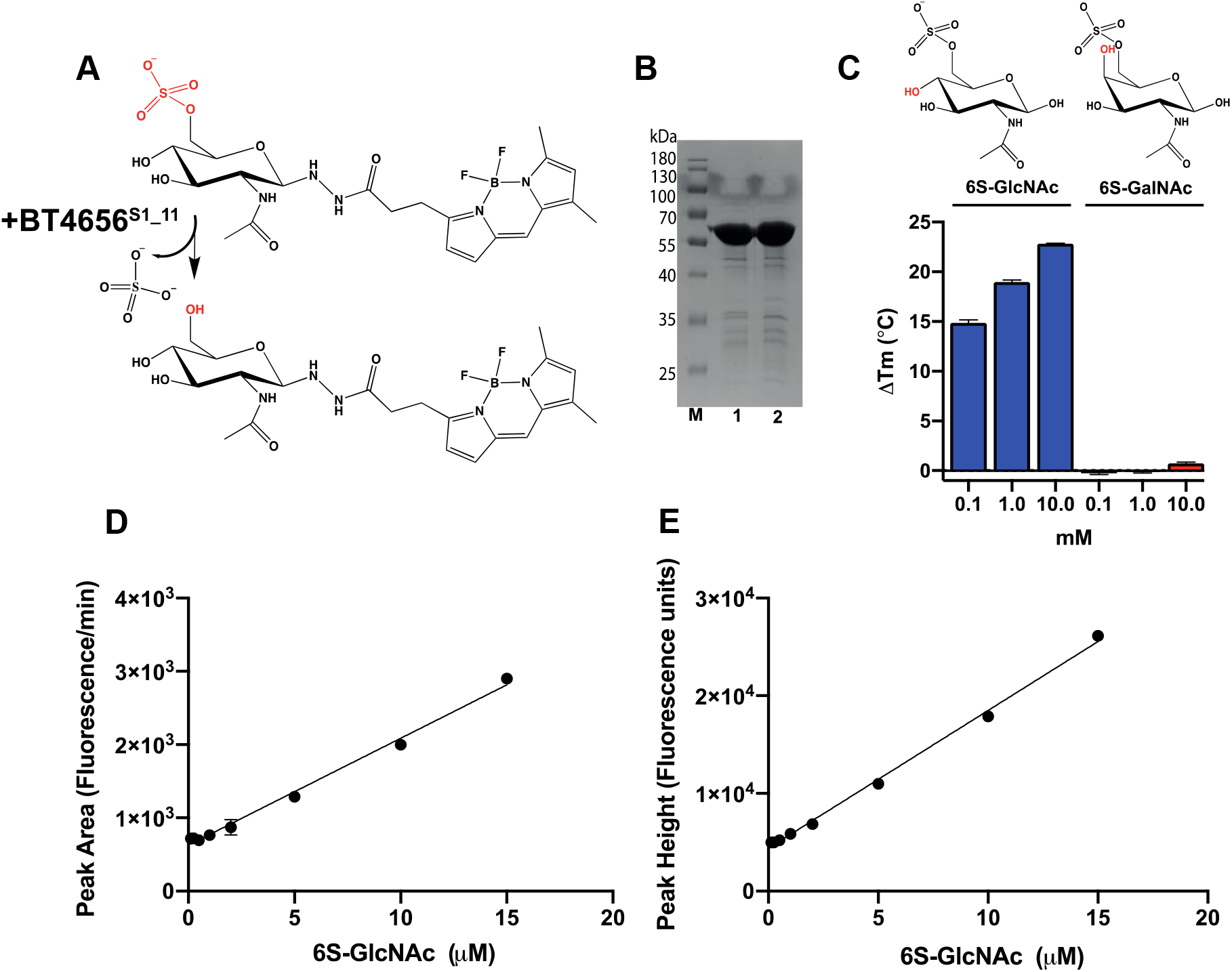
Substrate and enzyme analysis, and concentration-dependent linear detection of BODIPY-labelled 6S-GlcNAc substrate using a high-throughput microfluidic assay. **A)** Structure of the BODIPY-FL hydrazide labelled substrate 6S-GlcNAc substrate, the ß anomer is depicted, but glycosylamines are also able to undergo mutarotation, and the structure of the product after action by BT4656^S1_11^. **B)** M is the protein ladder and lanes 1 and 2 contain catalytically-active Hexa His-tagged BT4656^S1_11^ (Ser77Cys) and the catalytically-inactive (wild-type) Hexa His-tagged BT4656^S1_11^ (Ser77) carbohydrate sulfatase, respectively, both migrating at ~62kDa. Both were purified to near homogeneity using cobalt based immobilised affinity chromatography. After dialysis, 8.5 μg of purified protein was separated by SDS-PAGE and stained with Coomassie Blue. **C)** Differential scanning fluorimetry was used to calculate midpoint denaturation temperature (Tm) values for the catalytically inactive wild-type form of BT4656^S1_11^ in the presence of its cognate substrate, 6S-*N*-acetyl-glucosamine, and the non-binding C4 epimer 6S-*N*-acetyl-galactosamine; **D)** LED-induced fluorescence of fluorescent substrate using EZ Reader II system. Peak area and **E)** peak height was plotted as a function of substrate concentration. Assays were conducted as described in the Methods.

**Figure 2.**
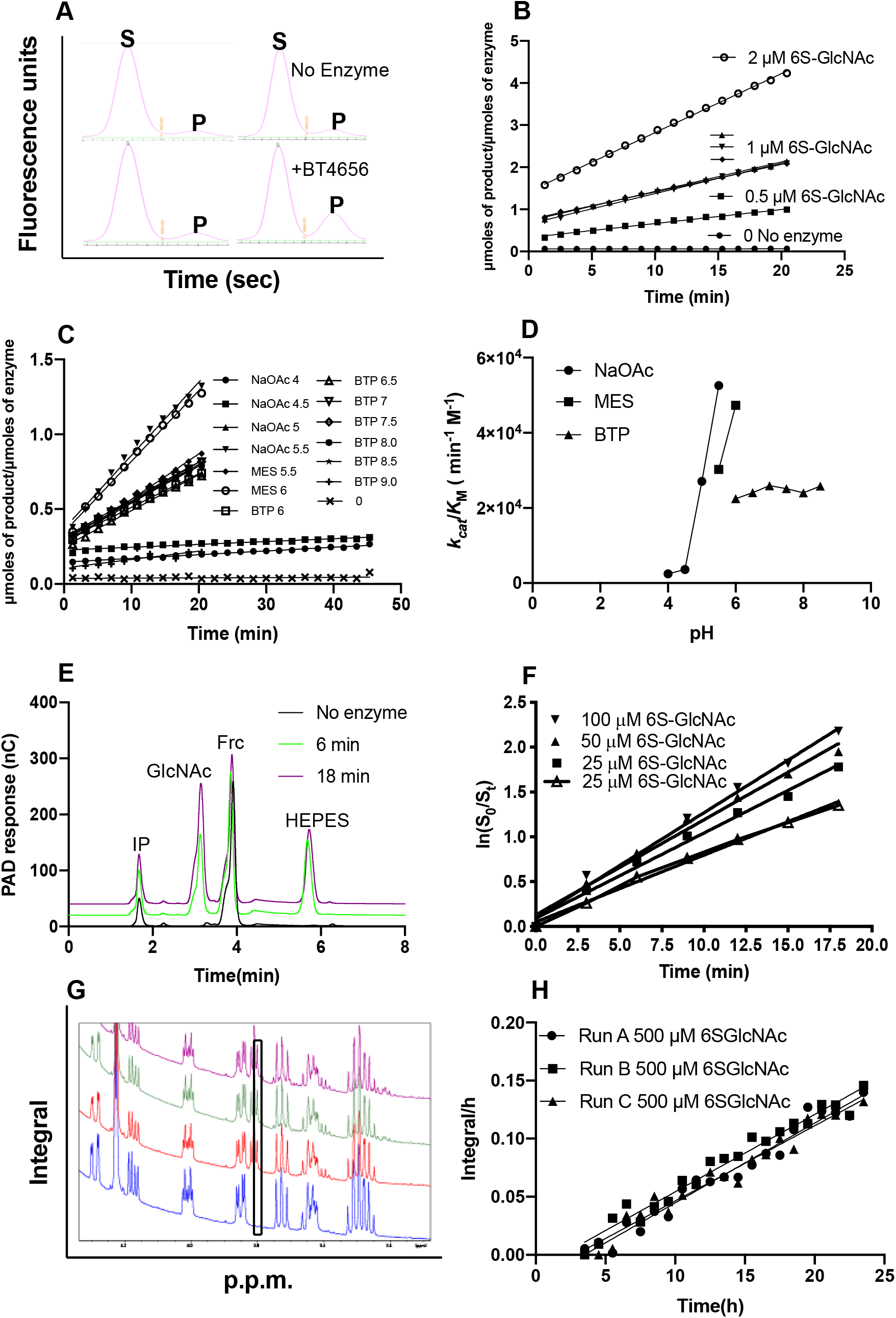
Side-by-side comparison of enzyme activity and kinetic data calculated using ca pillary electrophoresis (CE) and microfluidics, HPAEC or NMR. **A)** Raw capillary electrophoresis data outputs generated by the Perkin Elmer EZ Reader II software. S indicates sulfated substrate peak and P desulfated product peak, a snapshot of an identical time point is shown in the presence and absence of the sulfatase; **B)** *k*_cat_/*K*_M_ determinations calculated using capillary electrophoresis coupled to fluorescence detection, using 100 nM BT4656^S1_11^; **C)** linear rates produced at a range of pH values to determine the pH optimum for BT4656 using 1 μM substrate and 350 nM BT4656^S1_11^; **D)** *k*_cat_/*K*_M_ determination produced from C plotted against pH; **E)** Raw data from HPAEC chromatograms, IP= injection peak, GlcNAc= N-Acetylglucosamine product produced by BT4656^S1_11^, Frc=Fructose used as an internal standard to enable accurate quantification between runs and HEPES indicates dialysis bu ffer ‘contamintion’;**F)** *k*_cat_/*K*_M_ determinations produced using HPAEC coupled to PAD using 800 nM BT4656^S1_11^; **G)** Raw integrals from NMR experiment using 500 μM substrate and 10 nM nM BT4656^S1_11^; **H)** Specific activity produced from raw NMR data presented in G, the black box indicates the appearance of an unsulfated O6 substrate, which increases with time.

Based on peak-sizes and relative migration properties, only ~7-8 % of the prepared synthetic substrate was un-sulfated prior to the assay **(Figure 2A)**. However, importantly, in the absence of enzyme, the amount of un-sulfated substrate remained constant over the course of the reaction, demonstrating its stability under these assay conditions. In contrast, incubation with the purified 6S sulfatase BT4656^S1_11^ led to an increase in the slower-migrating desulfated product and a relative decrease in the faster-migrating sulfated substrate peak (**Figure 2A**). Indeed, repeated sampling of substrate and product peak height ratios allowed de-sulfation to be monitored in the same experiment, essentially in real-time, over the duration of the assay. Significantly, the total assay time, including experimental set-up and data generation, was less than one hour, and the procedure utilized very low amounts of both enzyme and substrate, leading at least an order-of-magnitude saving when time, assay and reagent costs are taken into account (see Table 1, and discussed below).

**Table 1.** Comparative values for *k*_cat_/*K*_M_ determined for purified sulfatase using three distinct analytical approaches. The table list the kinetic values and the corresponding amount of resources required in terms of time and components.* Indicates value is a specific activity and a *k*_cat_/*K*_M_ determination. All experiments represent technical triplicates.

|  | CE-Fluorescence | HPAEC-PAD | NMR | Ratio<br>HPAEC:<br>CE | Ratio<br>NMR:<br>CE |
| --- | --- | --- | --- | --- | --- |
| **$k_{cat}/K_M$ ($\text{min}^{-1} \text{M}^{-1}$)** | $(6.4 \pm 0.1) \times 10^4$ | $(2.0 \pm 0.4) \times 10^5$ | $^*(3.3 \pm 0.06) \times 10^2$ | ~3:1 | NA |
| **Time taken for $k_{cat}/K_M$ determination** | <1h | ~40h | ~12h | ~40:1 | ~12:1 |
| **Mass of substrate used** | 0.08 $\mu\text{g}$ | 48 $\mu\text{g}$ | 339 $\mu\text{g}$ | ~600:1 | ~4238:1 |
| **Mass of enzyme used** | 1.56 $\mu\text{g}$ | 19.38 $\mu\text{g}$ | 1.35 $\mu\text{g}$ | ~12:1 | ~0.87:1 |
| **Reaction volume** | 80 $\mu\text{l}$ | 1000 $\mu\text{l}$ | 700 $\mu\text{l}$ | 12.5:1 | 8.75:1 |
| **Time points recorded** | 16 | 6 | 24 | - | - |

### Kinetic analysis of carbohydrate de-sulfation

We next determined the appropriate quantity of enzyme required to enable measurement of initial rates of carbohydrate de-sulfation by the sulfatase. An enzyme concentration of 100 nM was selected for CE-based desulfation assays, since this generated ~20 % substrate hydrolysis ~20-40 minutes after assay initiation in a total assay volume of 80 μl. Once optimum experimental parameters had been established, the kinetic parameters of the BT4656^S1_11^ sulfatase could be determined rapidly using our fluorescence coupled-CE assay procedure. In these experiments, we determined the *k*_cat_/*K*_M_ for BT4656^S1_11^ to be 6.4 x 10^4^ min^-1^ M^-1^ using fluorescent sulfated substrate at concentrations of 0.5, 1 and 2 μM **(Figure 2 B and Table 1)**. The high throughput, 384 well plate format of the assay also enabled us to establish that BT4656^S1_11^ possesses a pH optimum of around pH 6.0 under these conditions (**Figure 2C, D**).

To compare our new CE-based protocol with previously established literature-standard carbohydrate de-sulfation assays, we measured BT4656^S1_11^ desulfation kinetics using HPAEC and NMR, the two ‘gold standard’ assays. High performance anion exchange chromatography (HPAEC) coupled with PAD, although relatively sensitive, lacks the enhanced sensitivity of the fluorescence-coupled microfluidic assay (which detects nM concentrations of desulfated carbohydrate product), and as such, *k*_cat_/*K*_M_ determinations could only be made using significantly higher substrate concentrations of 25, 50 and 100 μM. The analytical HPAEC approach is based on substrate depletion and accurate measurements require >70 % hydrolysis[35]. Therefore, to ensure high rates of substrate turnover, 800 nM final concentration of the enzyme was employed and the assay was conducted discontinuously over ~20 mins in a 1 ml volume prior to analysis. In comparison to our CE based method, in which multiple experimental conditions and relative desulfation can be evaluated simultaneously, each discontinuous HPAEC experiment requires ~1 hour to perform, thus, a single *k*_cat_/*K*_M_ determination with 7 samples measured over 20 mins has a total run time of ~7-8 hours, making the overall data collection for *k*_cat_/*K*_M_ determination using three different substrate concentrations ~40 hours. This is an under-estimate of the real time needed, however, because it excludes initial optimisation experiments, which are trivial using the present CE-based procedure, and which render HPAEC highly inefficient compared to the new method, when using identical preparations of enzyme and substrate. As an example, the CE-based pH analysis presented in Figure 1D, which took ~1 hour, would have taken ~100 hours using HPAEC and would require prohibitively large quantities of substrate to reach essentially the same conclusion. Furthermore, whereas CE distinguishes both sulfated and de-sulfated GlcNAc within the same reaction, detection by pulsed amperometry only detects product formation, which is a major potential issue when performing substrate depletion assays, especially if end-products have an effect on the rate of enzymatic activity. To resolve this confounding issue, the reaction must therefore be driven to completion and then the percentage of substrate remaining at each time point retrospectively back-calculated from the product formed.

### NMR-based analysis of sulfatase activity

Similar to our new CE-based assay, analysis by NMR spectroscopy permits direct ‘visualisation’ of both the 6S-GlcNAc substrate formed and the GlcNAc product, and the kinetic rates can also be measured continuously. However, in marked contrast, NMR requires more substantial quantities for each experiment, e.g. 500 μM substrate was employed in a 700 μl volume to generate the data shown in **Table 1**. For expensive, or technically-challenging synthetic substrates (which is the case for most carbohydrate substrates, such as sulfated Lewis antigens), this becomes restrictive for NMR analysis. To perform the experiment in triplicate using an auto-sampler, the experiment must also be performed over several hours to allow sufficient time for sample loading and unloading. As such, 10 nM of BT4656^S1_11^ was selected, in order to maintain a low rate of substrate turnover. In addition, owing to the use of a single substrate concentration, which has to be relatively high due to sensitivity issues **(Table 1)**, only the specific activity could be determined **(Figure G, H)**, rather than a *k*_cat_/*K*_M_ value, which was readily calculated using both CE and HPEAC methods. Other factors to consider with standard NMR desulfation methods are the need for costed access to a spectrometer and the expertise required to run and analyse the experiment. The choice of buffer must also be appropriate for use in NMR; chemicals such as HEPES, which have heterocyclic rings, cause signal overlap with those from 6S-GlcNAc, and so were avoided. No such restrictions exist using CE-based sulfatase assays.

The CE analytical platform allowed rapid (minutes), facile *k*_cat_/*K*_M_ and pH calculations for BT4656^S1_11^ using low μM amounts of substrate and protein in a small (80 μl) reaction volume, an invaluable advantage when either is limited (see **Table 1** for a side-by-side comparison of data obtained using each technique). Furthermore, following addition of enzyme, the assays can be measured continuously in a high throughput (384 well plate) format, allowing for simultaneous analysis of a variety of experimental conditions. One caveat of CE is that the substrate needs to be modified by fluorescent labelling at a site distal to the modification. However, virtually all carbohydrate active enzymes characterised to date function by performing catalysis at the non-reducing end of the molecule, making them particularly amenable to analysis using this approach, since labelling is achieved readily at their reducing ends (see below). A final consideration is that the commercial LED used in this study operates in a fairly narrow, non-adjustable, 450-490 nm excitation range, restricting the fluorophores that can be employed but, which could be remedied easily through the relatively straightforward introduction of alternative LEDs or by employing an array detector.

### Enzyme inhibition studies using CE-based desulfation detection method

The recent development of CE-based assays for tyrosylprotein sulfatases allowed us to discover several new classes of PAPS-competitive small molecule inhibitor, including compounds previously identified as kinase inhibitors [31]. We next analysed the potential inhibitory effects of inorganic anions on BT4656^S1_11^ by including sulfate (the product of the sulfatase reaction) alongside structurally-similar phosphate and vanadate anions in our CE sulfatase enzyme assays. As expected, both sulfate and phosphate anions displayed significant inhibition only at high (10 mM) concentrations, whilst more potent inhibition was observed using lower concentrations (1 mM) of vanadate ions. This enabled us to calculate a *Ki* value for vanadate of 116.7 μM **(Figure 3A, B).**

**Figure 3.**
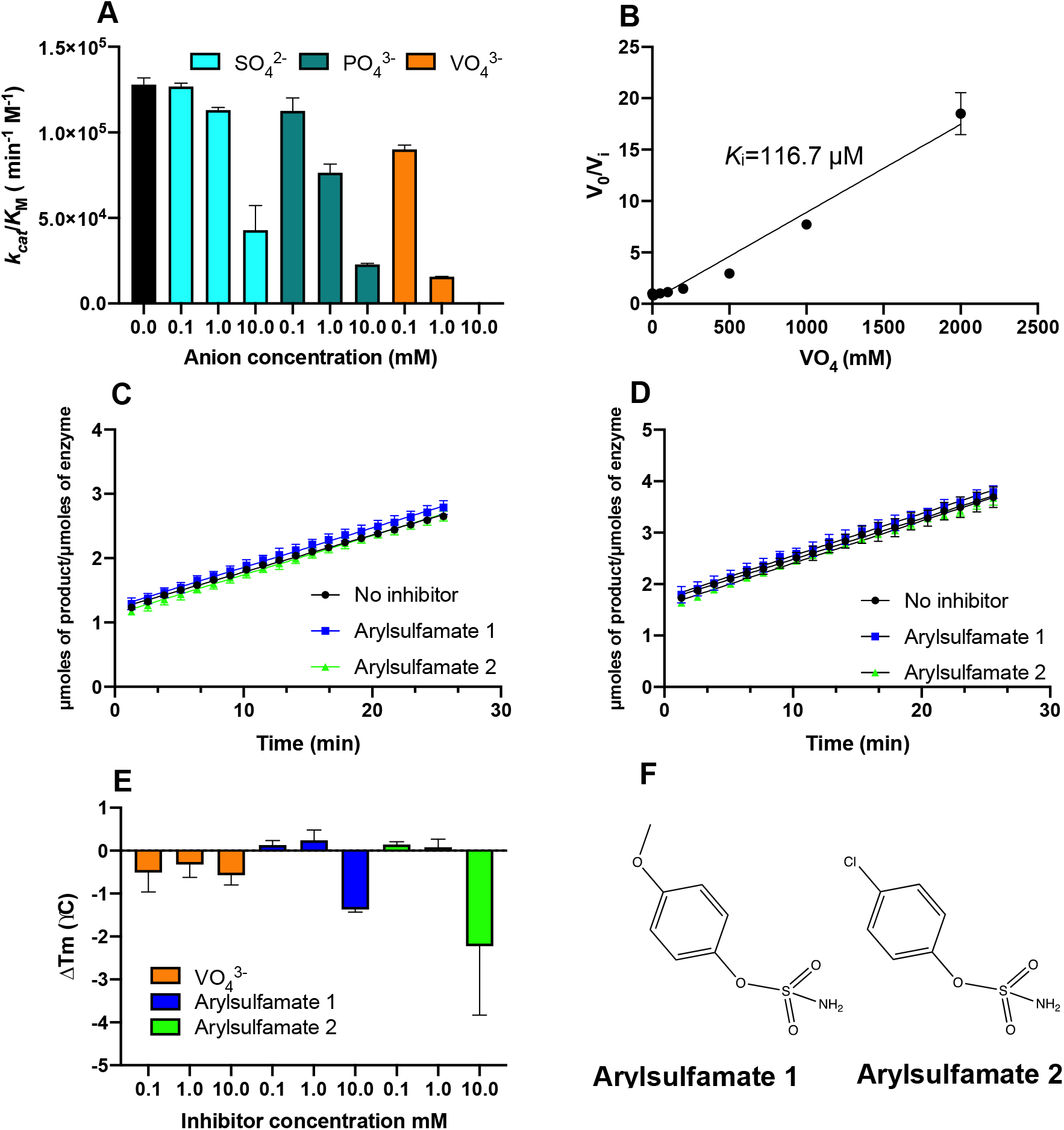
Evaluation of inhibition by arylsulfamates and enzyme product mimics. **A)** Inhibitory measurements for the product sulfate (SO_4_^2-^, cyan bars), and the putative sulfate product-mimics phosphate (PO_4_^3-^, teal bars) and vanadate (VO_4_^3-^, orange bars), black bars are controls lacking any addition; **B)** *K*_i_ determination for vanadate using the equation 1/*K*_i_= ((V_0_/V_i_)-1)/ [I]. All data are n=3; **C)** Inhibition measurements using 2 μM substrate, 100 nM BT4656^S1_11^ and 10 mM of either compound. 10 % DMSO was present in all reactions, including control incubations; **D)** Inhibitory measurements using 2 μM substrate, 100 nM BT4656^S1_11^ after incubating BT4656^S1_11^ with 5 mM arylsulfamates for 16 h. 5 % DMSO was present in all enzyme reactions; with each experiment performed in technical triplicate; **E)** Analysis of vanadate (VO_4_^3-^, orange bars) and arylsulfamates (blue and green bars) binding assessed by differential scanning fluorimetry (DSF); **F)** Structure of two arylsulfamate compounds analysed as potential inhibitors of BT4656^S1_11^.

We next analyzed inhibition of BT4656^S1_11^ by the arylsulfamate class of active-site steroid sulfatase inhibitors. Arylsulfamates have been shown to act on a broad range of sulfatases, but were originally developed as steroid sulfatase inhibitors[36] Surprisingly, the arylsulfamates investigated here (**Figure 3C, D, E, F**) did not lead to any marked inhibition of BT4656^S1_11^, either when measured via continuous assay in the presence of 10 mM compound (**Figure 3C**) or after overnight incubation of the sulfatase with 5 mM of the respective compounds (**Figure 3D**). DSF analysis of purified sulfatase binding to arylsulfamates demonstrated that they did not thermally stabilise BT4656^S1_11^, consistent with their lack of enzymatic inhibition. There was only a mild destabilising effect, relative to the very marked stabilization observed for the natural 6S-*N*-acetyl-glucosamine substrate, at the highest arylsulfamate concentration tested, (**Figure 1C**), but this had no effect on the catalytic rate measured (**Figure 3C, D**). In contrast, BT4656^S1_11^ was not significantly stabilized or destabilized by vanadate, despite its ability to cause significant inhibition at μM concentrations. This lack of thermal protection suggests that vanadate-binding does not result in a marked conformational change on the sulfatase. The ineffectiveness of the tested arylsulfamates towards BT4656^S1_11^ is likely due to this family of sulfatases having evolved to target polar carbohydrate substrates, rendering the enzyme incompatible with the binding of aryl moieties, the hydrophobic substrates of the aryl sulfatases. Significant advances have been made in the design of inhibitors targeting steroid sulfatases involved in hormone dependent cancers and some are in Phase II clinical trials[37]. There are, however, currently no targeted, effective inhibitors of carbohydrate sulfatases despite their emergence as novel targets for disease intervention[25, 38, 39]. Taken together, our data demonstrate that CE can be used to study inhibition of a carbohydrate sulfatase in real-time using small amounts of sample, which will make this assay ideal for medium throughput analysis using compound libraries in a 384 well plate.

### Generation of Fluorescently-labelled Glycan substrate for CE assays

As described above, the fluorescent tag, BODIPY-FL hydrazide (λ505/513 and ε 80,000 M^-1^cm^-1^) was selected for the labelling of the reducing end of the carbohydrate substrates to generate a fluorescent 6S-GlcNAc-BOPIDY substrate that can be enzymatically desulfated by the sulfatase and detected in real-time alongside any product formed. The chemistry of coupling to the reducing end aldehyde group of sugars can be explained by a number of potential mechanisms which depend on the conditions and the nature of the labelled sugar. These different routes, some of which are shown in Figure 4A and B, can potentially lead to subtly different products including, in the case of reductive amination (an additional popular route for labelling sugars), open-ring products (initially forming imines (Schiff’s bases) or, in their reduced form, open chain amines). However, since we wished to avoid formation of fixed open ring structures to better mimic the natural carbohydrate substrate, a labelling method was selected based on a published procedure for the formation of glycosylamines [40], which allow closed ring forms to predominate.

**Figure 4.**
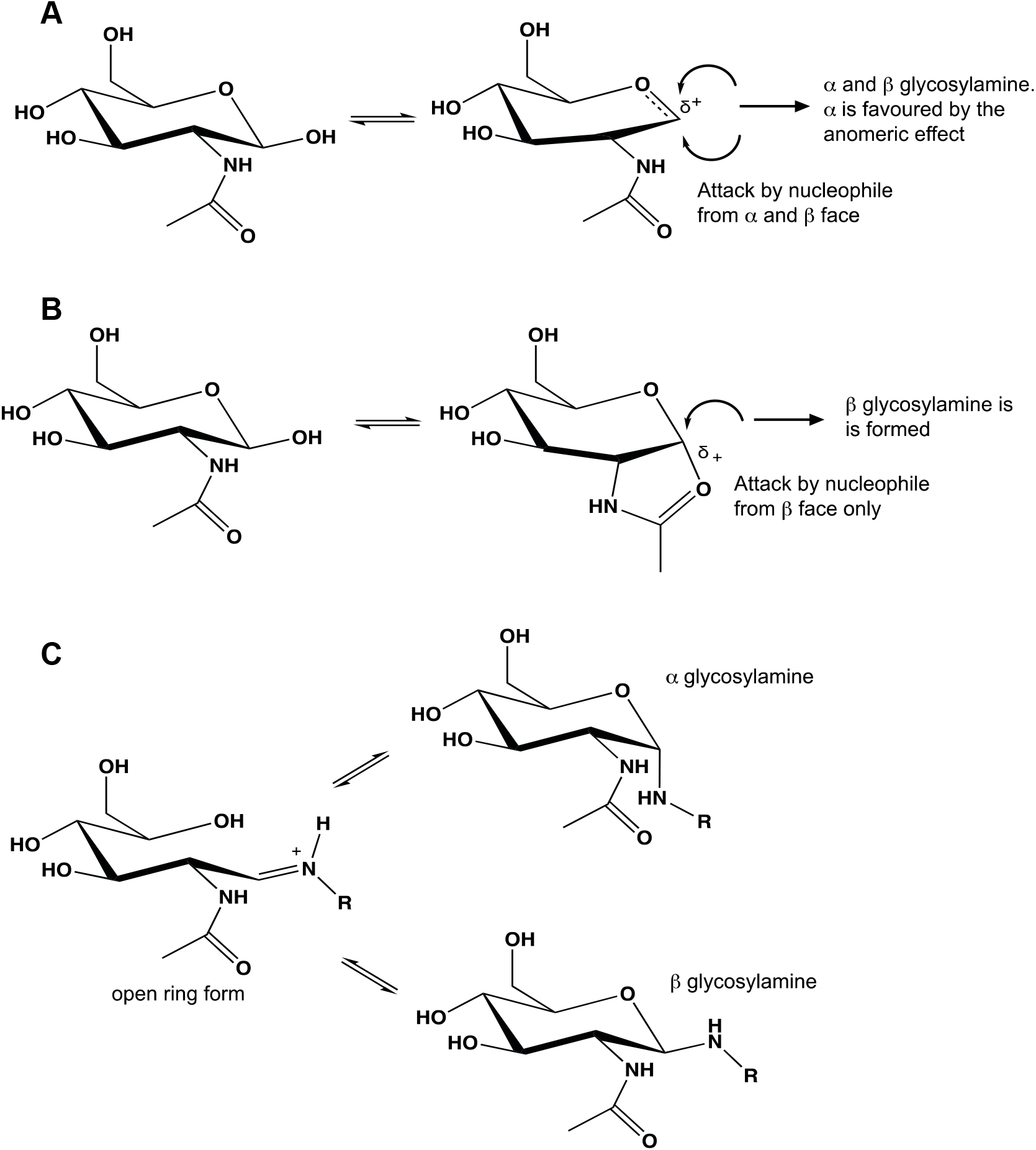
Labelling mechanism and anomeric configuration of products. The observed (approx. 1:1) mixture of the glycosylamine products of the reaction between 2-amino benzoic acid and *N*-acetylated-d-hexosamines can be envisaged as arising from two possible mechanisms; **A)** Proceeding via an oxocarbenium ion intermediate produces both possible anomers, but the α form is expected to dominate owing to the anomeric effect, although this is a weaker effect for nitrogen than for oxygen; **B)** Proceeding through an oxazoline intermediate resulting exclusively in the β anomer; **C)** The glycosylamine product, however, may be subject to mutarotation via an open chain form allowing equilibration of the α and ß anomers. R= BODIPY-FL hydrazide or 2-amino-benzoic acid (2-AA).

To further explore the structure of the products formed by this method, an analogous coupling was conducted between both GalNAc and GlcNAc and 2-amino-benzoic acid (2-AA) and the products were analysed by NMR spectroscopy. This confirmed formation of the closed ring glycosylamine products in both anomeric forms (**Figure 5**). The reaction between the reducing end of the sugar and an amine-containing label (or hydrazide, which exhibit similar reactivity in polar solvents [41]) could proceed conceivably by several mechanisms including formation of an oxacarbenium (glycosyl) cation or, as in the present case of *N*-acetylated hexosamines, via formation of an oxazoline form (oxazolinium ion). Each of these mechanisms (**Figure 4A, B**) has slightly different outcomes in terms of the anomeric products formed. Unfortunately, the ability of the generated *N*-linked glycosides to undergo mutarotation allows equilibration between both α and β anomers [42] prevents the mechanism of formation from being deduced unequivocally **(Figure 4C)**. Nevertheless, we believe that this facile workflow, in which a synthetic fluorescent substrate can be both desulfated by recombinant bacterial sulfatase, and readily detected and quantified after capilliary electrophoresis, is an important advance for the carbohydrate sulfatase field.

**Figure 5.**
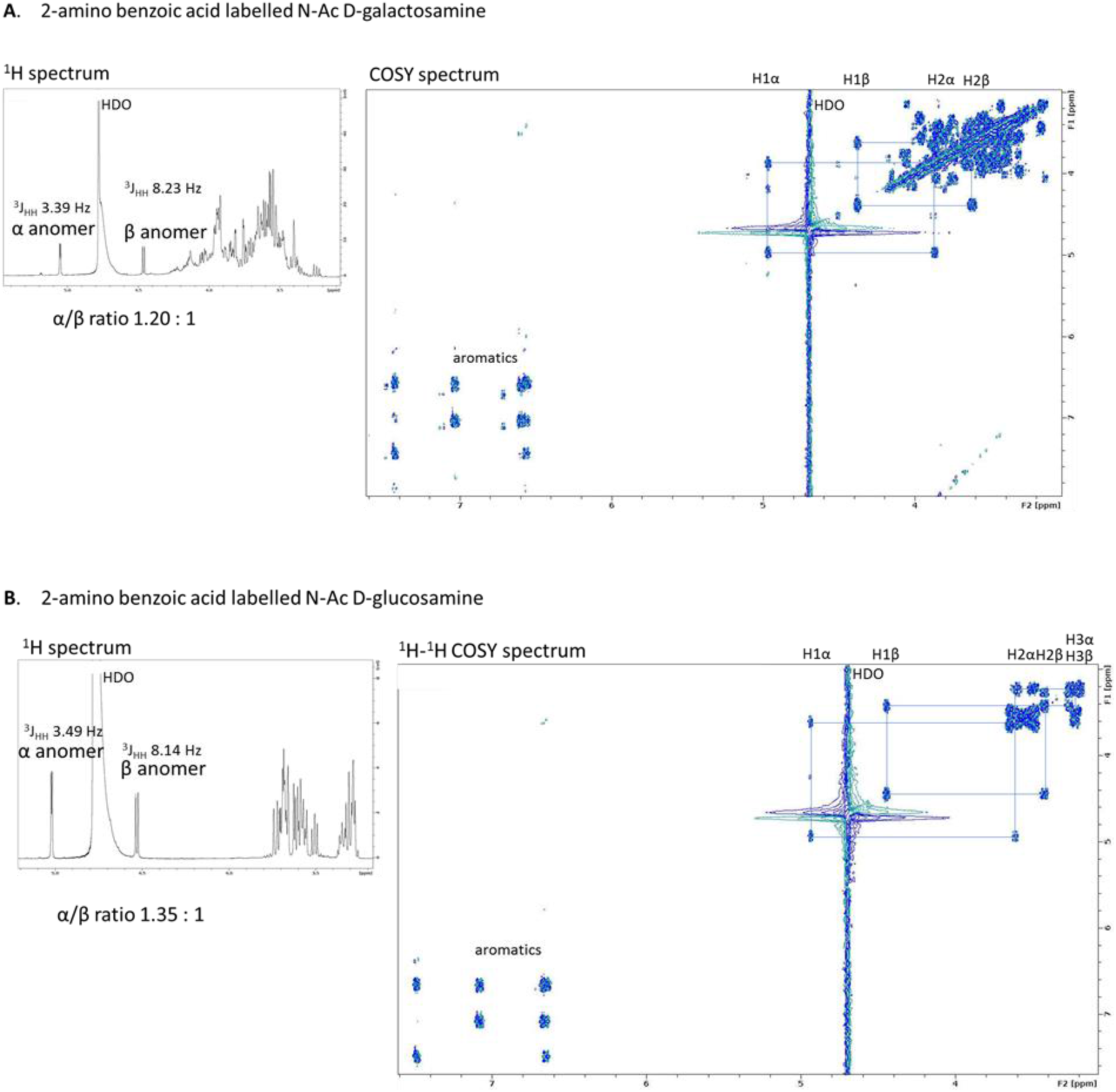
NMR analysis of glycosylamine linked derivatives. **A)** NMR spectra of 2-amino benzoic acid derivative of *N*-acetyl-d-galactosamine (left) ^1^H NMR spectrum of product showing characteristic α and β anomeric signals (right) ^1^H-^1^H COSY NMR spectrum with partial assignments shown. **B)** NMR spectra of 2-amino benzoic acid derivative of *N*-acetyl-d-glucosamine (left) ^1^H NMR spectrum of product showing characteristic α and β anomeric signals (right) ^1^H-^1^H COSY NMR spectrum with partial assignments shown.

## Conclusion

In this study, we show for the first time that capillary electrophoresis coupled to fluorescence-based product detection offers a rapid, facile and high-throughput method for quantitative carbohydrate sulfatase analysis. Enzymatic desulfation of a fluorescent sulfated carbohydrate substrate by a purified recombinant sulfatase requires less time and material than more conventional methods of carbohydrate analysis, such as HPAEC and NMR. Moreover, our method can be used to generate kinetic and enzyme inhibition data that might be extended to other carbohydrate sulfatases, and for the analysis of other sulfated and/or phosphorylated glycans, in the future. Finally, our fluorescent 6S-GlcNAc substrate could also be used in screens for assay optimization amongst other sulfatases, or for the discovery of inhibitors of a variety of carbohydrate sulfatases, including those implicated in human diseases.

## Abbreviations

GAG: Glycosaminoglycan
6S-GlcNAc: 6S-*N*-acetyl-d-glucosamine
6S-GalNAc: 6S-*N*-acetyl-d-galactosamine
HPAEC: High performance anion exchange chromatography
DSF: Differential Scanning Fluorimetry
CE: capillary electrophoresis
FG: Formylglycine
S1: formylglycine containing family of sulfatases

## Data availability statement

The datasets generated during and/or analysed during the current study are available from the corresponding authors on reasonable request.

## COMPETING INTERESTS

There are no perceived conflicts of interest from any authors.

## FUNDING SUPPORT

AC was supported by the academy of medical sciences/Wellcome Trust through the springboard grant SBF005\1065 163470 awarded. JAL and EAY acknowledge support from the BBSRC project BB/M011186/1. DPB and PE acknowledge generous funding from BBSRC grants BB/S018514/1 and BB/N021703/1 and North West Cancer Research (NWCR) grants CR1097 and CR1208.

## Methods

### Recombinant Protein Production

Catalytically active recombinant BT4656^S1_11^ (in which Ser 77 was mutated to Cys, to permit prokaryotic conversion of the thiol to an aldehyde and the generation of the formylglycine nucleophile), or catalytically-inactive BT4656^S1_11^ (in which Ser 77 is intact but prevents formation of formylglycine during expression in *E.coli*) were expressed in *Escherichia coli* strain TUNER (Novagen), and cultured to mid-exponential phase in LB supplemented with 50 μg/mL kanamycin at 37 °C and 180 rpm. Cells were then cooled to 16 °C, and recombinant gene expression was induced by the addition of 0.1 mM isopropyl β-D-1-thiogalactopyranoside; cells were cultured for another 16 h at 16 °C and 180 rpm. The cells were then centrifuged at 5,000 × g and resuspended in 20 mM HEPES, pH 7.4, with 500 mM NaCl before being sonicated on ice. Recombinant protein was then purified by immobilized metal ion affinity chromatography using a cobalt-based matrix (Talon, Clontech) and eluted with 100 mM imidazole. Proteins were then analysed by SDS-PAGE gel and appropriately pure fractions dialysed into 10 mM HEPES pH 7.0 with 150 mM NaCl. Protein concentrations were determined by measuring absorbance at 280 nm using the molar extinction coefficient calculated by ProtParam on the ExPasy server (web.expasy.org/protparam/).

### A novel microfuidic-based desulfation assay

6S-*N*-acetyl-d-glucosamine was labelled at its reducing end with BODIPY which has a maximal emission absorbance of ~503nm, which can be detected by the EZ Reader II platform [32] *via* LED-induced fluorescence. Non-radioactive microfluidic mobility shift carbohydrate sulfation assays were optimised in solution with a 12-sipper chip coated with CR8 reagent and performed using a PerkinElmer EZ Reader II system employing EDTA-based separation buffer and real-time kinetic evaluation of substrate desulfation. Pressure and voltage settings were adjusted manually (1.8 psi, upstream voltage: 2250 V, downstream voltage: 500 V) to afford optimal separation in the reaction mixture of the sulfated substrate and unsulfated glycan product, with a sample (sip) time of 0.2 s, and total assay times appropriate for the experiment. Individual desulfation assays were carried out at 28°C and were pre-assembled in a 384-well plate in a volume of 80 μl in the presence of substrate concentrations between 0.5 and 20 μM with 100 mM Bis-Tris-Propane or Tris, dependent on the pH required, and 150 mM NaCl, 0.02% (v/v) Brij-35 and 5 mM CaCl_2_. The amount of de-sulfation was directly calculated in real-time using EZ Reader II software by measuring the sulfated carbohydrate: unsulfated carbohydrate ratio at each time-point during the assay. The activity of sulfatase enzymes was quantified in ‘kinetic mode’ by monitoring the amount of unsulfated glycan generated over the assay time, relative to control assay with no enzyme; with sulfate elimination from the substrate limited to ~20% to prevent loss assay linearity via substrate depletion. *k*_cat_/*K*_M_ values, using the equation V_0_=(V_max_/*K*_M_)/S, were determined by linear regression analysis with GraphPad Prism software. Substrate concentrations were halved and doubled to check linearity of the rates ensuring substrate concentrations were significantly <*K*_M_.

### HPAEC-based desulfation assays

The ability of BT4656^6S-sulf^ to remove O6-linked sulfate from GlcNAc6S was monitored by the production of GlcNAc detected by high-performance anion exchange chromatography (HPAEC) with pulsed amperometric detection using a carbohydrate standard quad waveform for electrochemical detection at a gold working electrode with an Ag/AgCl pH reference electrode and a Carbopac PA-100 guard and analytical column (Dionex; ThermoFisher). Assays were performed in 1 ml volumes. After first removing a zero time point sample, enzyme was added, then 110 μl volumes were removed, and boiled to destroy enzyme activity, at periodic time intervals over 18 minutes (Figure 2E, F). Samples were then centrifuged and subjected to HPAEC. GlcNAc was separated using an isocratic gradient ran over 20 minutes with 100 mM NaOH, the column was then stripped and washed with 10 min runs of 100 mM NaOH plus 500 mM Sodium acetate then 10 mins with 500 mM NaOH before being ran back into 100 mM NaOH ready for the next sample. A flow rate of 1.0 ml/min was used throughout.

### NMR-based detection of glycan desulfation

NMR experiments were designed to monitor the desulfation of 6S-*N*-acetyl-d-glucosamine and were conducted in D2O with 50 mM sodium phosphate, pH 7.0, supplemented with 150 mM NaCl at 25°C on a 600MHz Bruker Avance II+ spectrometer, fitted with a TCI CryoProbe. 1D proton and 2D ^1^H, ^1^H-COSY spectra were measured using standard pulse sequences provided by the manufacturer. Spectra were processed and analysed using TopSpin 3.4A and TopSpin 4.0 software (Bruker). *N*-acetyl-d-glucosamine integrals were recorded indirectly for desulfation of C(6)H_2_-OSO_3_ to C(6)H_2_-OH using the emerging C(4) peak as a proxy, within the region 3.395 to 3.442ppm, referenced to the combined C(3) peaks for *N*-Acetyl-d-glucosamine and 6S-*N*-acetyl-d-glucosamine in the region 3.679 to 3.762ppm.

### Differential scanning fluorimetry

Thermal shift/stability assays (TSAs) were performed using a StepOnePlus Real-Time PCR machine (LifeTechnologies) and SYPRO-Orange dye (emission maximum 570 nm, Invitrogen) with thermal ramping between 20 and 95°C in 0.3°C step intervals per data point to induce denaturation of purified, folded wild-type (Ser 77) BT4656^S1_11^ in the presence or absence of carbohydrates, anions, and arylsulfamates. The melting temperature (Tm) corresponding to the midpoint for the protein unfolding transition was calculated by fitting the sigmoidal melt curve to the Boltzmann equation using GraphPad Prism, with R^2^ values of ≥0.99, as described in[43]. Data points after the fluorescence intensity maximum were excluded from the fitting. Changes in the unfolding transition temperature compared with the control curve (ΔTm) were calculated for each ligand. A positive ΔTm value indicates that the ligand stabilises the protein from thermal denaturation, and confirms binding to the protein. All TSA experiments were conducted using a final protein concentration of 5 μM in 100 mM Bis-Tris-Propane (BTP), pH 7.0, and 150 mM NaCl, supplemented with the appropriate ligand concentration. Three independent assays were performed for each protein and protein ligand combination.

### Carbohydrate labelling

To a dried sample of the sugar to be labelled (1.5 μmoles) in a screw-capped microfuge tube (1.5 mL), was added BODIPY-FL hydrazide (0.48 mg, 1.65 μmoles (1.1 eq., ThermoFisher) dissolved in 0.5 mL anhydrous methanol (Sigma)). The reaction was conducted at 65°C (overnight) in a heat block. Upon completion, the reaction was cooled and a sample of the product separated by thin layer chromatography on aluminium-backed silica (Millipore) developed in methanol (2 ascents; labelled 6S-GlcNAc product, orange, Rf 0.77 and free label, BODIPY-FL hydrazide, also orange; Rf 0.60) A sample was recovered by scrapping the silica from the plate and the product was extracted in methanol (0.5 mL x 3), followed by filtration. The purified product was recovered by evaporating the solvent (rotary evaporator) and was stored dry at −20°C in the dark.

## Notes

### Competing Interest Statement

The authors have declared no competing interest.

